# Translocation of effector proteins into plant cells by the flax rust pathogen *Melampsora lini*

**DOI:** 10.1101/2024.12.01.625101

**Authors:** Xiaoxiao Zhang, Ann-Maree Catanzariti, Gregory J. Lawrence, Pamela H. P. Gan, David A. Jones, Peter N. Dodds, John P. Rathjen

## Abstract

During infection, rust fungi secrete effector proteins into host plant cells from haustoria to aid their colonisation. How rust effectors are secreted from the haustorium and delivered into the cytoplasm of host cells remains poorly understood. We used an *Agrobacterium*-mediated transformation procedure to generate stable transgenic flax rust strains expressing the effectors AvrM and AvrP123 fused to yellow fluorescent protein (YFP). We showed that both AvrM-YFP and AvrP123-YFP fusion proteins were secreted by the fungus into the extrahaustorial matrix (EHMx); however only AvrM-YFP was delivered into host cells, triggering a typical resistance phenotype in plants carrying the corresponding resistance (*R)* gene *M*. The signal peptide of AvrM was sufficient to direct YFP secretion into the EHMx; however, delivery into the host cell required a larger 105 amino acid N-terminal fragment of AvrM. These results indicate that translocation of this protein into the host cell from the EHMx is a separate process from secretion into the EHMx and requires a signal present in AvrM between amino acids 34 and 105. This is in contrast to previous observations of AvrM localisation after transient expression in plants, highlighting the necessity for analysis in the natural infection system.

## Introduction

Plant pathogens secrete effector proteins that contribute to virulence by suppressing host immunity, amongst other functions (He *et al*., 2020). Broadly speaking, effectors may be secreted into apoplastic spaces or translocated into the cytoplasm of host cells. Cytoplasmic effectors can be recognised by host resistance (R) proteins of the nucleotide-binding leucine-rich repeat (NLR) sub-class, leading to effector-triggered immunity (ETI) (Dodds & Rathjen, 2010). ETI is often accompanied by the hypersensitive response (HR) including cell death and restriction of the pathogen to the infection site.

Delivery of effectors is best understood in pathogenic bacteria, which use a type-III secretion system to deposit effectors directly into the host cytoplasm (Alfano & Collmer, 2004). Type-III effectors are produced in the bacterial cytosol and are unfolded by chaperone proteins followed by direct injection into the host cell via a needle-like pilus. Eukaryotic pathogens including oomycetes and fungi seem to have evolved analogous effector delivery systems independently in various lineages (Lo Presti & Kahmann, 2017; Bozkurt & Kamoun, 2020). Oomycetes such as the potato blight pathogen *Phytophthora infestans* deliver effectors both to the apoplast and the cytoplasm of host cells (Wang *et al*., 2017). Both types of effectors carry classical signal peptides for secretion via the endoplasmic reticulum co-translational pathway. However, *Phytophthora* apoplastic effectors are secreted through the conventional Golgi-mediated pathway that can be disrupted by the inhibitor brefeldin A (BFA), whereas cytoplasmic effectors seem to be secreted through a BFA-insensitive pathway (Wang *et al*., 2017). The cytoplasmic oomycete effectors include the RXLR sub-class, characterised by a conserved RxLR-dEER sequence motif that follows the N-terminal signal peptide (Whisson *et al*., 2007). The RXLR motif appears to be required for protein delivery into the plant cells through an as yet unknown mechanism, possibly diverting the secreted proteins into the Golgi-independent secretory pathway (Wang *et al*., 2017). The RxLR region appears to be cleaved from the effector protein prior to secretion (Wawra *et al*., 2017). The effectors of plant pathogenic fungi do not contain recognisable RxLR motifs and the mechanisms of delivery remain poorly characterised. Two distinct secretion pathways were also identified in the ascomycete fungus *Magnaporthe oryzae.* The host-delivered *M. oryzae* effectors are secreted through a biotrophic interface complex (BIC) into the extrahyphal space and the cytoplasm of rice host cells via a Golgi-independent pathway, while extracellular effectors are secreted from invasive hyphae through the conventional Golgi-mediated pathway (Giraldo *et al*., 2013). Recent independent studies in both *P. infestans* (Wang *et al*., 2023b) and *M. oryzae* (Oliveira-Garcia *et al*., 2023) suggested that uptake of cytoplasmic effectors into the plant cells are facilitated by clathrin-mediated endocytosis (CME). However, the CME pathway may not be the only route of effector entry into plant cells used by fungal and oomycete pathogens (Wang *et al*., 2023a). For example the basidiomycete *Ustilago maydis* also delivers effectors into host cells but does not form BIC-like structures, suggesting an alternative means of delivery (Tanaka *et al*., 2015). Indeed, a protein complex containing at least seven *U. maydis* secreted proteins including two intrinsic membrane proteins accumulates at the cell surface and is required for virulence, suggesting a possible function as a translocon for delivery of pathogen effectors (Ludwig *et al*., 2021).

The rust fungi are a large family of basidiomycete pathogens that feed solely on live host cells, causing significant yield losses to important agricultural crops (Aime *et al*., 2017; Figueroa *et al*., 2020). Infection typically starts from spore germination on the surface of the infected leaf, followed by development of an infection peg that enters the leaf through a stoma (Kobayashi *et al*., 1994; Garnica *et al*., 2014). The fungus then develops a substomatal vesicle and infection hyphae that ramify throughout the apoplastic spaces of the leaf. After contacting leaf mesophyll cells, the hyphae differentiate into haustorial mother cells which form a specialized structure that penetrates the plant cell wall and enlarges between the cell wall and the host plasma membrane, developing into the haustorium (Kobayashi *et al*., 1994; Garnica *et al*., 2014). The haustorium is thought to be the primary site of nutrient acquisition from the host and enables further development of the fungus including massive spore production. The infection cycle is completed when the fungus forms uredospores that erupt through the leaf epidermis in characteristic pustules (Garnica *et al*., 2014).

A number of effector proteins have been identified from the flax rust fungus *Melampsora lini* through genetic studies of its interaction with its host plant flax (*Linum usitatissimum*). These effectors trigger host immunity upon recognition by corresponding flax NLR proteins and are therefore termed avirulence (Avr) proteins. *Avr* genes have been cloned from six independent loci in flax including *AvrL567*, *AvrL2*, *AvrM*, *AvrM14*, *AvrP123* and *AvrP4*, all of which encode proteins that are expressed in haustoria and contain a canonical N-terminal signal peptide (SP) for protein secretion (Dodds *et al*., 2004; Catanzariti *et al*., 2006; Barrett *et al*., 2009; Anderson *et al*., 2016). Recognition by intracellular host NLR proteins indicates that these Avr proteins are delivered into plant cells upon secretion. Similarly, six Avr proteins identified from the wheat stem rust fungus *Puccinia graminis* f. sp. *tritici* are expressed in haustoria and recognised inside host cells directly by the corresponding host NLR proteins (Chen *et al*., 2017; Salcedo *et al*., 2017; Upadhyaya *et al*., 2021; Arndell *et al*., 2024) or indirectly by a host NLR through a helper R protein (Chen *et al*., 2024).

Direct recognition of rust fungus Avr proteins by physical interaction with cytosolic R proteins has been shown in nine cases (Dodds *et al*., 2006; Catanzariti *et al*., 2010; Chen *et al*., 2017; Salcedo *et al*., 2017; Deng *et al*., 2022; Chen *et al*., 2023; Arndell *et al*., 2024; Chen *et al*., 2024; Kim *et al*., 2024; Shen *et al*., 2024), further supporting their delivery into host cells. Immunolocalisation of AvrM from flax rust as well as the RTP1 effector from bean rust (*Uromyces fabae*) detected these proteins inside host cells during infection (Kemen *et al*., 2005; Rafiqi *et al*., 2010). Transcriptome analysis of several species of rust fungi has identified a large number of genes that are expressed specifically in haustoria and encode predicted SPs; however it is not clear which of them are host-delivered and which are retained in the apoplastic space (Duplessis *et al*., 2011; Garnica *et al*., 2013; Cuomo *et al*., 2017; Miller *et al*., 2018; Schwessinger *et al*., 2018; Li *et al*., 2019). In general, effectors from rust fungi lack a typical RxLR motif and no BIC-like structures have been described, suggesting that an effector delivery mechanism has evolved independently in these fungi.

The rust haustorium is surrounded by two membrane layers (Coffey *et al*., 1972; Littlefield & Bracker, 1972; Szabo & Bushnell, 2001). The first is the fungal haustorial plasma membrane, which is surrounded by the haustorial cell wall and the extrahaustorial matrix (EHMx). The second is the extrahaustorial membrane (EHM), which is continuous with but differentiated from the plant plasma membrane (PM). The EHMx is a gel-like layer rich in carbohydrates and is sealed from the plant apoplastic space by a neckband at the site of cell wall penetration. Secreted effectors are expected to be delivered to the EHMx via signal peptide-mediated secretion, but how they move from the EHMx into host cells is not known. To further investigate effector protein translocation during rust infection, we employed stable transgenic *M. lini* strains expressing effector:fluorescent protein fusions to follow their localisation in fungal and host structures during infection. These transgenic rust strains enabled direct observation of effector protein translocation in infected plant tissues using confocal microscopy. We showed that the *M. lini* effector AvrM was secreted into the EHMx and delivered into host cells by the pathogen, resulting in ETI in flax lines containing the corresponding *R* genes. The signal peptide of AvrM was sufficient for protein secretion into the EHMx but delivery of the protein into plant cells required additional N-terminal AvrM sequences.

## Results

### Delivery of AvrM-YFP into flax cytoplasm by transgenic *M. lini*

To investigate rust effector delivery, we generated stable transgenic *M. lini* strains expressing the effector protein AvrM fused to YFP to enable live-cell imaging using confocal microscopy. Two independent AvrM-YFP transgenic rust strains, tAvrM-YFP-S1 and tAvrM-YFP-S2, were examined. These were propagated by growth on the susceptible flax *cv*. Hoshangabad on which they were fully virulent. We first investigated whether the AvrM-YFP protein was recognised by the host *R* gene *M*. The two transgenic rust strains were independently inoculated onto the susceptible flax line *cv*. Hoshangabad and three *M*-containing flax lines: *cv*. Dakota, and backcross lines containing the *M* gene in either the Hoshangabad (Dakota x Hoshangabad) or Bison (Dakota x Bison) backgrounds (Fig 1A), and observed for the development of rust pustules. The non-transformed parental *M. lini* strain CH5F2-96 that does not carry *AvrM* was able to grow and sporulate on *cv*. Hoshangabad and all three *M*-containing flax lines, indicating a virulent phenotype. Both tAvrM-YFP-S1 and tAvrM-YFP-S2 were able to grow on *cv*. Hoshangabad, however produced typical avirulence phenotypes on *M*-containing flax lines with no sporulation and small hypersenstive flecks observed on the infected leaves. Consistent with this, chitin measurements showed that growth of tAvrM-YFP-S1 and tAvrM-YFP-S2 were inhibited in the *M*-containing *cv*. Dakota x Hoshangabad compared to that of CH5F2-96 (Fig 1B). This suggested that the AvrM-YFP fusion protein was properly expressed and recognised by the cytosolic M resistance protein to confer avirulence. Expression of the AvrM-YFP fusion protein was confirmed by immunoblotting with anti-GFP antibodies of leaf protein extracts from flax *cv*. Hoshangabad infected with tAvrM-YFP-S1 and tAvrM-YFP-S2 but was not detected in CH5F2-96 infected samples (Fig. 1C).

**Fig 1.**
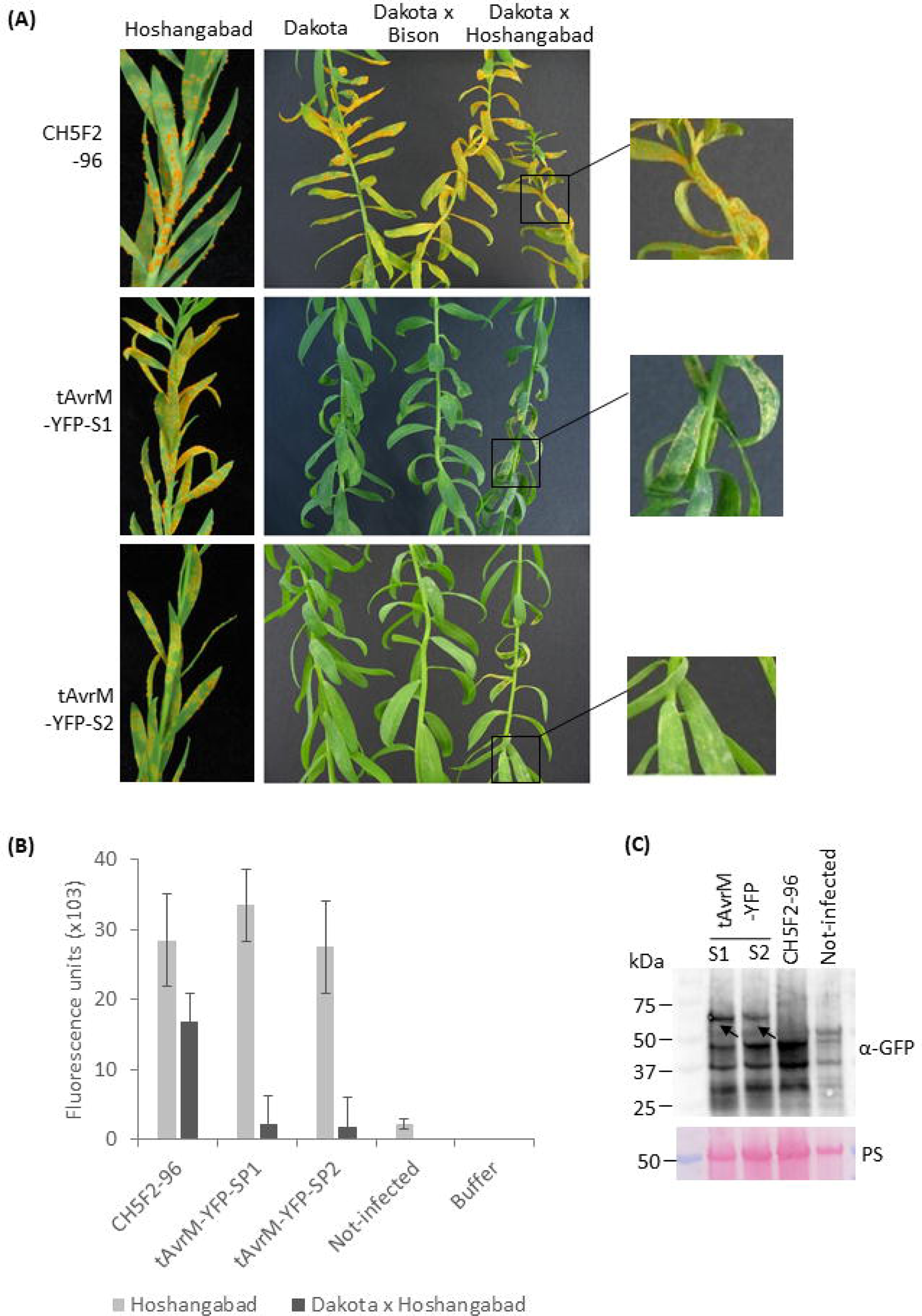
Disease phenotypes and protein production of AvrM-YFP in flax leaves infected with transgenic *M. lini* strains. **(A)** Disease phenotypes of the parental strain CH5F2-96 and transgenic *M. lini* tAvrM-YFP-S1 and tAvrM-YFP-S2 on susceptible flax *cv.* Hoshangabad and *M*-containing flax *cv.* Dakota and Hoshangabad or Bison backcross derivatives (Dakota x Bison or Dakota x Hoshangabad). Zoomed regions are outlined by frames and showed in the right panel to show different phenotypes. Yellowing on the leaves corresponds to sporulating pustules and white hypersensitive flecks correspond to localised cell death. The plants were imaged at 10 dpi. **(B)** Quantification of *M. lini* growth on flax *cv*. Dakota x Hoshangabad and *cv*. Hoshangabad by the wheat germ agglutinin chitin (WAC) assay. Each column represents the average fluorescence obtained from three replicate samples. The error bars represent the standard deviation. **(C)** Immunoblotting analysis of AvrM-YFP protein expression in leaves of flax *cv*. Hoshangabad infected with tAvrM-YFP-S1, tAvrM-YFP-S2 or CH5F2-96 using anti-GFP antibodies. The arrows indicate bands at the expected molecular weight of AvrM-YFP. Protein loading is indicated by Ponceau staining (PS) of the Rubisco large subunit.

We next investigated the growth of the transgenic rust strains and accumulation of AvrM-YFP by light and confocal microscopy at 3 or 4 days post inoculation (dpi). These time points were chosen because the expression of *M. lini Avr* genes including *AvrM* and *AvrP123* was previously shown to peak at 2-5 dpi (Wu *et al*., 2019). At 3 dpi, hyphae were observed primarily at stomata between the guard cells (Fig 2A and 2B), with very few haustoria-like structures identified. YFP-specific fluorescence was not detected in the 3 dpi samples. Non-YFP fluorescence detected in fungal hyphae and substomatal vesicles showed a spectral profile that was different from a typical YFP fluorescence profile (Fig S1A). At 4 dpi, significant hyphal growth was evident and haustoria could be observed within flax cells (Fig 2C-2F). In addition, YFP-specific fluorescence (Fig 2C, 2E and S1B) was detected, indicating accumulation of the AvrM-YFP fusion protein.

**Fig 2.**
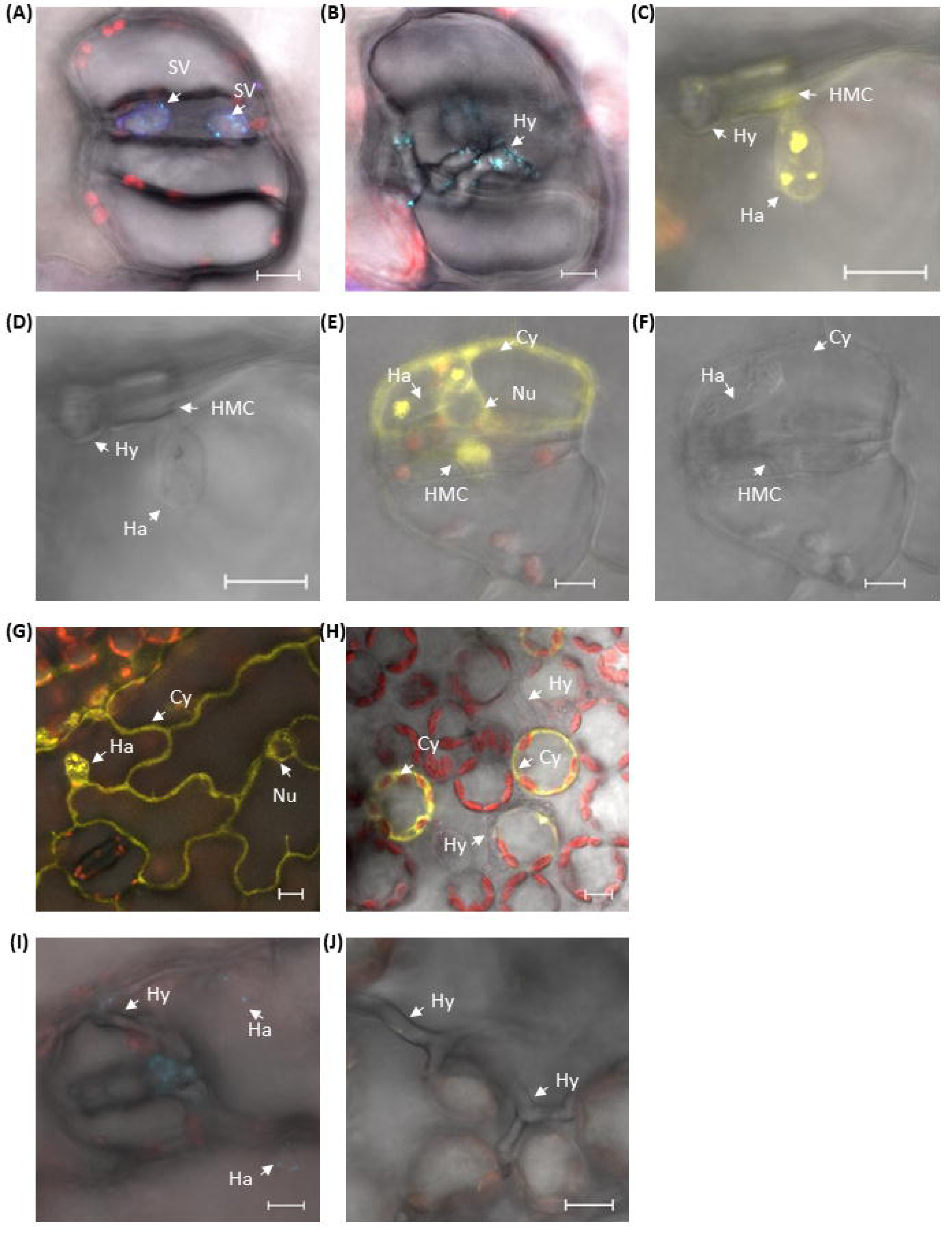
AvrM-YFP localisation in flax leaves infected with transgenic *M. lini* strains. **(A and B)** Linear unmixing confocal fluorescence microscopy images showing infection by tAvrM-YFP-S1 at 3 dpi. The spectral images were unmixed and merged with bright field images. Chloroplast autofluorescence is shown in red; undefined fluorescence emissions are shown in blue and cyan. Representative images show flax stomata and guard cells with fungal substomatal vesicles (SV) in (a) and hyphae (Hy) in (B). **(C)** Merged confocal image showing AvrM-YFP localisation at 4 dpi. YFP-specific fluorescence is shown in yellow. **(D)** Bright field microscopic image of (C) showing infection by tAvrM-YFP-S1 at 4 dpi. **(E)** Merged confocal image as in (C). **(F)** Bright field microscopic image of (E). **(G and H)** Representative confocal images showing infection by tAvrM-YFP-S1 in flax epidermal cells and mesophyll cells respectively. **(I and J)** Representative confocal images showing infection by the non-transgenic *M. lini* strain CH5F2-96. SV, substomatal vesicles; Hy, hyphae; Ha, haustorium; HMC, haustorial mother cells; Cy, flax cytoplasm; Nu, flax nucleus. Scale bars indicate 10 µm.

YFP fluorescence in *M. lini* tAvrM-YFP strains was observed early in haustorial development, both in haustorial mother cells (HMC) and young haustoria (Fig. 2C and 2E). Maturing haustoria expanded and branched into typical multilobed structures, expressing YFP fluorescence within haustoria and at the haustorial periphery (Fig 2E). This latter observation may be consistent with secretion of AvrM-YFP into the extrahaustorial matrix (EHMx). YFP fluorescence was further observed in the cytoplasm of infected host cells that contained a haustorium, but not in uninfected cells (Fig 2E, 2G and 2H), which is consistent with previous immunolocalisation studies of AvrM during infection (Rafiqi *et al*., 2010). YFP fluorescence was detected in cells of different types that contained haustoria, including guard cells (Fig. 2E), epidermal cells (Fig. 2G) and mesophyll cells (Fig. 2H), but was absent from host cell nuclei (Fig. 2E and 2G). Nuclear exclusion of AvrM was also observed previously in immunolocalisation studies and after transient *in planta* expression of AvrM-YFP (Ve *et al*., 2013). In contrast, YFP fluorescence was not observed in plant samples infected with the untransformed parental strain CH5F2-96 at 4 dpi (Fig 2I and 2J). Furthermore, in flax cells transiently expressing the red fluorescent protein (RFP) in the cytosol, AvrM-YFP fluorescence from transgenic *M. lini* infections partially co-localized with RFP at the haustorial periphery (Fig 3A and 3B) and in the infected cell cytosol (Fig 3A and 3C). YFP fluorescence but not RFP was observed within haustoria (Fig 3A and 3B). Again, no YFP fluorescence was observed in plant tissue infected with the non-transformed *M. lini* strain CH5F2-96 (Fig 3D - 3F). RFP fluorescence was observed in the flax cytoplasm and surrounding the haustorial body but was not observed inside the haustorium, resulting in a ring-shape fluorescence structure. In flax cells not infected with *M. lini,* RFP fluorescence was observed in the cytoplasm (Fig 3G). Overall, these observations indicated that AvrM-YFP was produced and secreted from *M. lini* haustoria likely into the EHMx and delivered into the cytoplasm of host cells.

**Fig. 3.**
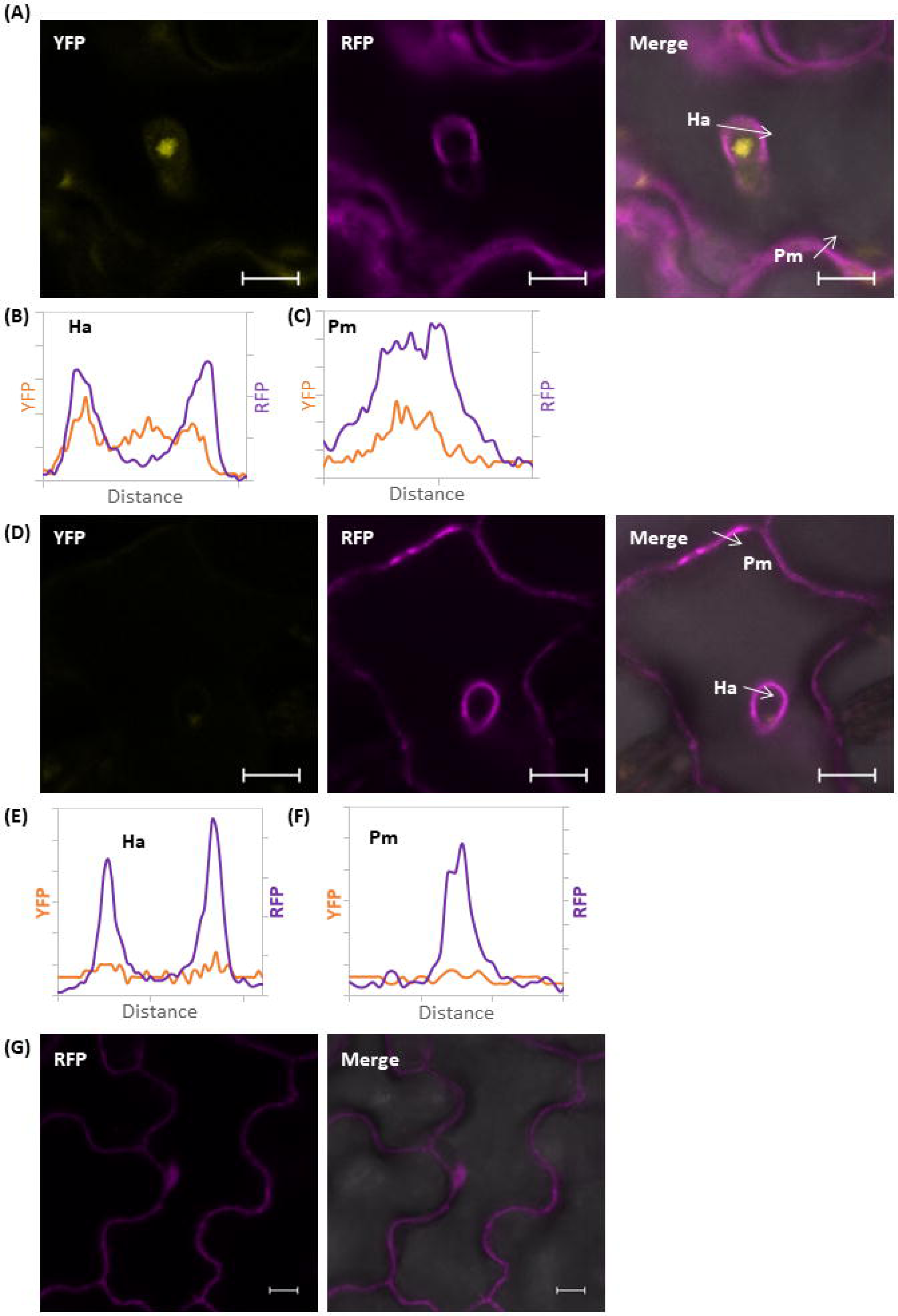
AvrM-YFP is delivered into the EHMx and host cytoplasm. **(A)** Confocal images showing infection by tAvrM-YFP-S1 in flax epidermal cells expressing cytosolic RFP. YFP-specific fluorescence appears in yellow; RFP fluorescence is shown in purple; chloroplast autofluorescence is shown in red. The arrows in the right panel indicate the lines used for the fluorescence intensity profiles in (B) and (C). Ha, haustorium; Pm, flax plasma membrane. **(B and C)** Fluorescence intensity profiles of (a). Each *x*-axis represents the distance from the end to the head of each arrow. **(D-F)** Confocal images and fluorescence intensity profiles showing infection by the parental strain CH5F2-96. **(G)** Confocal image showing RFP fluorescence in the absence of the fungus. Scale bars indicate 10 µm.

Focal accumulation of YFP fluorescence was also observed within the haustorial body, which often contained 1 to 3 small fluorescent bodies of 1-5 µm in size (Fig 2C, 2E and 2G). Figure 4A and 4B shows a close up of an expanded mature haustoria (which we defined as those which had secreted AvrM-YFP into the area surrounding the haustorium) showing YFP fluorescence gathering into two unknown structures, each of which was about 5 µm in diameter (Fig 4A). The two structures were surrounded by small unidentified dots emitting non-YFP fluorescence. The two structures and dots were partially visible in the brightfield (Fig 4B). In another less developed haustorium (Fig 4C & 4D), in which AvrM-YFP accumulated to a lower level and was not observed at the haustorial periphery, the non-YFP fluorescence foci and to a lesser extent YFP fluorescence clustered in two similar structures within the haustorium which were visible in the brightfield (Fig 4D). The size and location of these structures suggested their association with the two nuclei present in haustoria (Coffey *et al*., 1972; Littlefield & Bracker, 1972). However, we were not able to detect DAPI staining of haustorial nuclei, presumably due to a lack of DAPI penetration into these structures since host cell nuclei were stained (data not shown).

**Fig 4.**
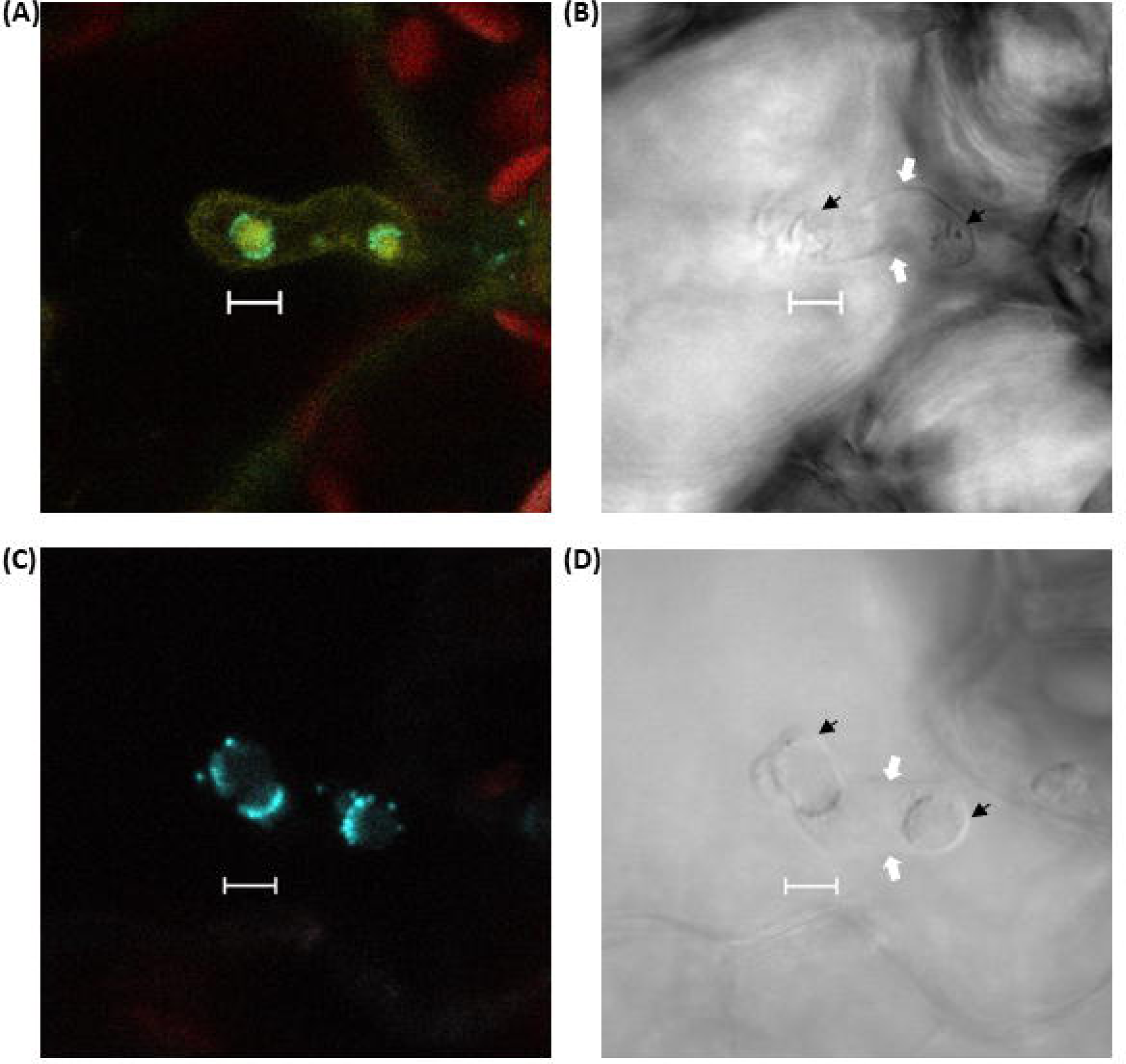
Accumulation of fluorescence within unknown haustorial substructures in transgenic *M. lini* expressing AvrM-YFP. **(A)** Representative linear unmixing of confocal fluorescence images showing AvrM-YFP fluorescence within a haustorium and at the haustorium periphery. YFP-specific fluorescence appears in yellow; chloroplast autofluorescence is shown in red; undefined fluorescence emissions are shown in cyan. **(B)** Bright field image of (A). The haustorial boundary is indicated by white arrows and the unknown fluorescent structures are indicated by black arrows. **(C)** Representative linear unmixing of confocal images showing AvrM-YFP localisation in the haustorium but not in EHMx coloured as (A). **(D)** Bright field image of (C) labelled as (B). Scale bars indicate 5 µm.

### AvrP123-YFP was not delivered into flax cytoplasm by transgenic *M. lini*

The *M. lini* effector AvrP123 is another haustorially-expressed secreted protein which is recognised by the intracellular flax R proteins P1, P2 and P3 (Catanzariti *et al*., 2006). To investigate the localisation of AvrP123 during infection, we transformed the *M. lini* strain CH5F2-74 which is virulent on flax lines expressing P2 with a construct expressing AvrP123-YFP under the control of its native promoter and terminator sequences. Two transgenic lines were examined; tAvrP123-YFP-S1 and tAvrP123-YFP-S4. Both strains gave similar infection levels to the virulent CH5-74 parent line when inoculated onto the susceptible flax line *cv*. Hoshangabad and the *P2*-containing flax line *cv* Abyssinian in the Hoshangabad background (Abyssinian x Hoshangabad) (Fig 5A). In contrast, the *M. lini* strain CH5 which carries *AvrP123* was avirulent on *cv*. Abyssinian x Hoshangabad. Chitin measurement showed that growth of CH5 but not CH5-74, tAvrP123-YFP-S1 or tAvrP123-YFP-S4 was inhibited in the *P2*-containing *cv.* Dakota x Hoshangabad (Fig 5B). Both transgenic lines expressed the fusion protein, as infection of *cv*. Hoshangabad with these strains followed by immunoblotting of leaf extracts with anti-GFP antibodies revealed bands corresponding to full-length AvrP123-YFP, whereas this band was absent from untransformed CH5F2-74 or an uninfected flax control (Fig 5C). This suggested that the AvrP123-YFP construct did not confer avirulence to the transgenic lines, despite expression of the fusion protein. We inoculated the tAvrP123-YFP-S1 strain onto flax *cv*. Hoshangabad to examine AvrP123-YFP localisation. YFP fluorescence was observed within haustoria and the EHMx but was absent from the host cytoplasm (Fig 5D and 5E). Similar YFP fluorescence patterns were observed in flax plants infected with tAvrP123-YFP-S4. Therefore, AvrP123-YFP was produced in both lines and secreted into the EHMx, but was not delivered into the plant cytoplasm. This explains the lack of an avirulence phenotype, and may be due to the YFP tag interfering with the correct delivery of the fusion protein.

**Fig 5.**
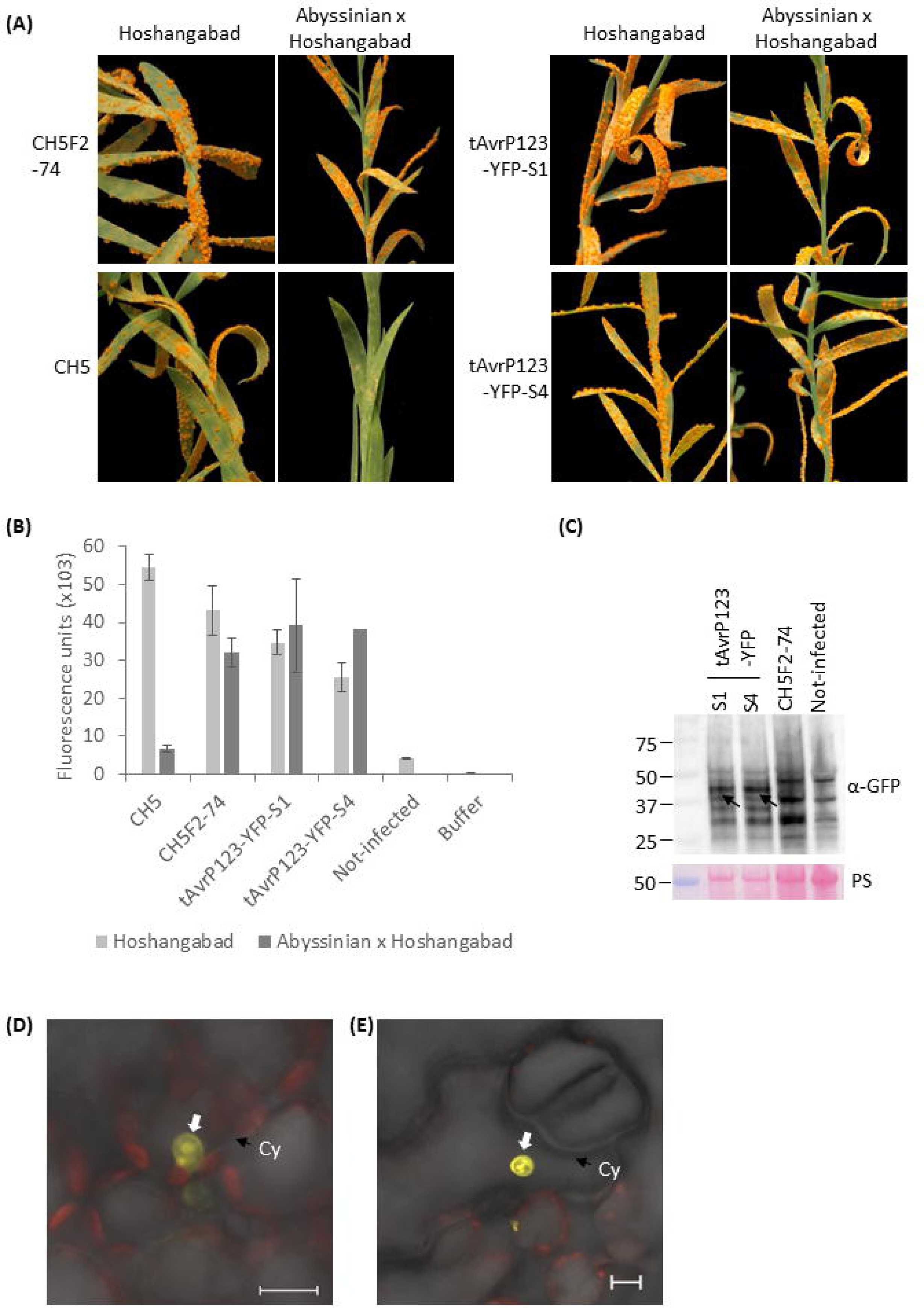
AvrP123-YFP localisation in flax leaves infected with transgenic *M. lini* strains. **(A)** Disease phenotypes observed on susceptible flax *cv*. Hoshangabad and *P2-*containing flax *cv.* Abyssinian x Hoshangabad after infection with the non-transgenic *M. lini* strain CH5 (contains *AvrP123*), parental strain CH5F2-74 (does not contain *AvrP123*) and transgenic strains tAvrP123-YFP-S1 and tAvrP123-YFP-S4. The images were taken at 14 dpi. Yellowing on the leaves corresponds to sporulating pustules. **(B)** Quantification of *M. lini* growth on flax *cv*. Hoshangabad and *cv*. Abyssinian x Hoshangabad by the wheat germ agglutinin chitin (WAC) assay. Each column represents the average fluorescence obtained from three replicate samples. The error bars represent the standard deviation. **(C)** Immunoblot analysis to detect AvrP123-YFP in flax leaf samples of *cv*. Hoshangabad infected with *M. lini* strains as shown. Arrows indicate the expected molecular weight of AvrP123-YFP. Protein loading is indicated by Ponceau staining (PS) of the Rubisco large subunit. **(D and E)** Representative confocal microscope image showing AvrP123-YFP localization in Hoshangabad leaves infected with tAvrP123-YFP-S1 at 4 dpi. YFP-specific fluorescence is shown in yellow and chloroplast autofluorescence in red. The white arrow indicates a haustorium. Cy, flax cytoplasm. Scale bars indicate 10 µm.

### An N-terminal fragment of AvrM is sufficient for YFP delivery into flax cells

To investigate features of the AvrM protein that enable delivery into host cells, we generated constructs containing various N-terminal fragments of AvrM fused to YFP, including the signal peptide alone (amino acid residues 1-34: SP-YFP) and two larger fragments of 105 and 156 amino acids (AvrM^1-105^-YFP and AvrM^1-156^-YFP; Table 1). Transgenic *M. lini* strains were examined for SP-YFP (1 line) and AvrM^1-105^-YFP (2 lines), but we did not obtain any transformants with the AvrM^1-156^-YFP construct. Fluorescence microscopy of susceptible flax infected with the transgenic line tSP-YFP detected YFP fluorescence in haustoria and the EHMx, however no signal was detected in the flax cytoplasm (Fig 6A). A further transgenic line containing a construct expressing the cyan fluorescent protein (CFP) alone (not fused to AvrM) from the *AvrM* promoter resulted in diffuse fluorescence throughout haustoria with no accumulation at the haustorial periphery or in host cytoplasm (Fig 6B). The difference in fluorescence profiles between these experiments suggests that SP-YFP was secreted into the EHMx whereas CFP which lacks a signal peptide was not. Thus, the AvrM SP fragment is sufficient for protein secretion into EHMx, but host delivery of the effector requires other features of the full-length protein. In contrast, expression of AvrM^1-105^-YFP fusion resulted in YFP fluorescence in haustoria, EHMx and the flax cytoplasm, similar to AvrM-YFP (Fig 6C and 6D). Notably, expression of AvrM^1-105^-YFP resulted in the accumulation of a strong YFP signal in the plant nucleus (Fig 6C), whereas the full length AvrM-YFP protein was excluded from the nucleus (Fig 2E and 2G). This is consistent with the smaller size of the AvrM^1-105^-YFP fusion protein allowing diffusion into the nucleus. In flax cells labelled with cytosolic RFP, the EHMx signals of SP-YFP (Fig 7A and 7B) and AvrM^1-105^-YFP (Fig 7C and 7D) partially co-localized with RFP at the haustorial periphery. In addition, AvrM^1-105^-YFP (Fig 7E) but not SP-YFP (Fig 7F) co-localized with RFP in the host cytosol. Both the SP-YFP and AvrM^1-105^-YFP proteins lack the C-terminal domain of AvrM recognised by M (Catanzariti *et al*., 2010) and were therefore not tested for avirulence phenotypes. Expression of the SP-YFP and AvrM^1-105^-YFP proteins by *M. lini* strains tAvrM^1-105^-YFP-S1, tAvrM^1-105^-YFP-S2 and tSP-YFP was confirmed by immunoblotting with an anti-GFP antibody (Fig 7G). Overall, the data indicate that the AvrM SP is sufficient for protein secretion into EHMx, but delivery of the effector requires an additional N-terminal region of AvrM between residues 34 and 105.

**Fig 6.**
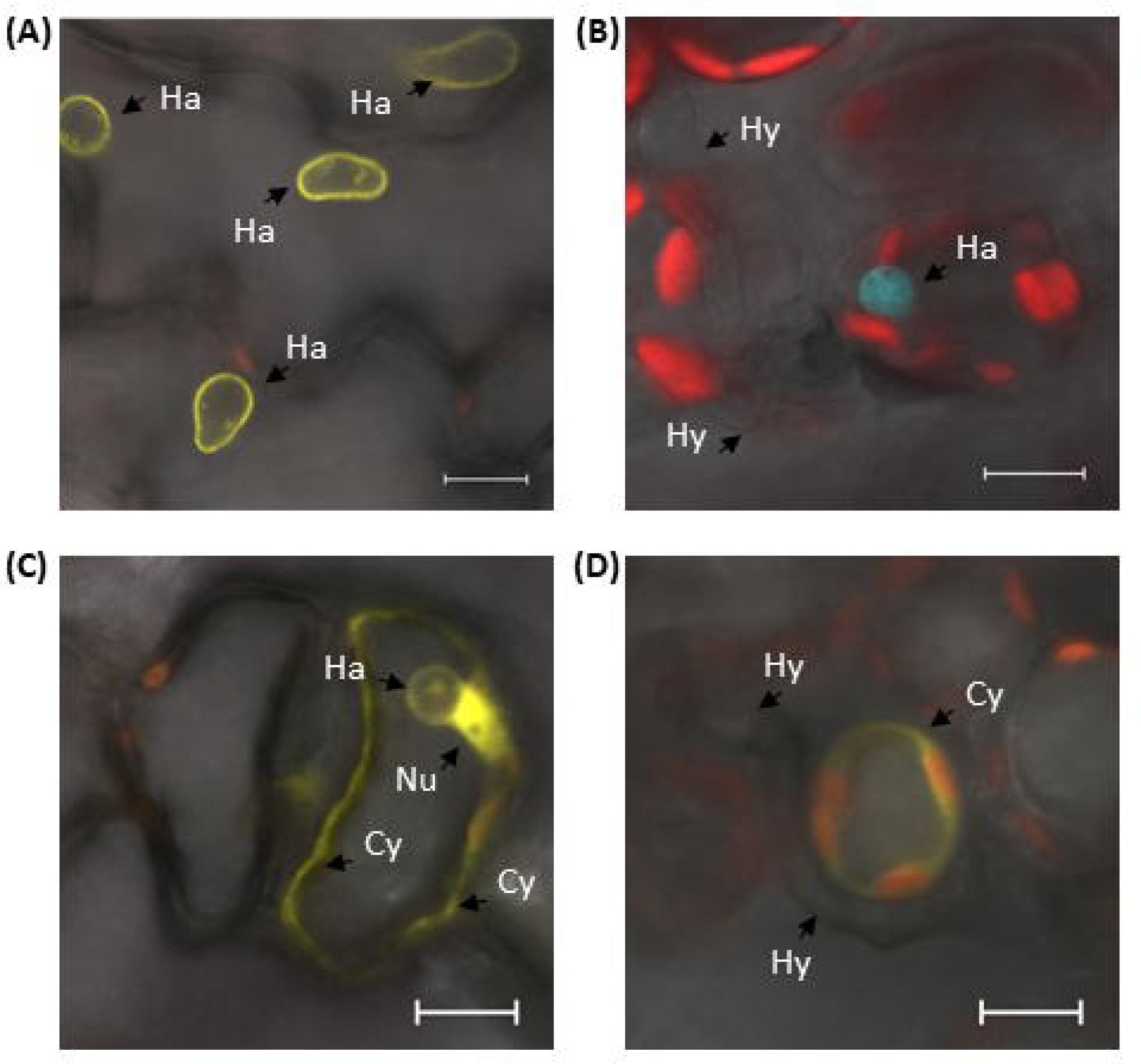
Localisation of SP-YFP and AvrM^1-105^-YFP proteins in flax leaves infected with the respective transgenic *M. lini* strains. **(A-D)** Representative confocal images showing localisation of SP-YFP (A), CFP (B) and AvrM^1-105^-YFP (C-D). YFP fluorescence is shown in yellow, chloroplast autofluorescence in red and CFP fluorescence in cyan. Scale bars indicate 10 µm.

**Fig 7.**
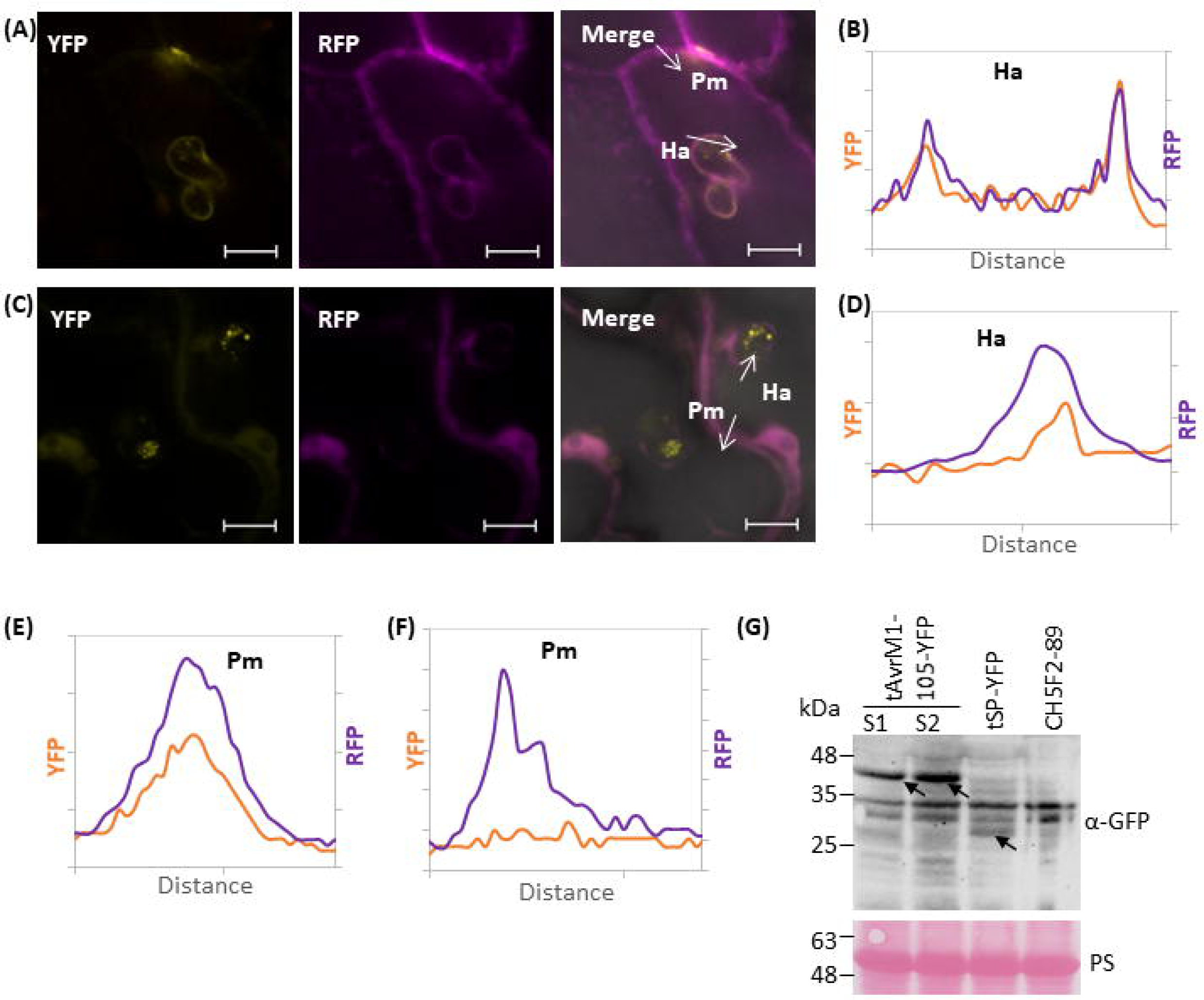
Localisation of SP-YFP and AvrM^1-105^-YFP proteins in flax leaves expressing cytosolic RFP. **(A)** Confocal images showing localisation of SP-YFP in flax epidermal cells expressing cytosolic RFP. The arrows indicate the lines used for the fluorescence intensity profiles across haustorium (Ha) or flax plasma membrane (Pm). Scale bars indicate 10 µm. **(B)** Fluorescence intensity profile of SP-YFP across Ha. **(C-D)** Confocal images and fluorescence intensity profile showing localisation of AvrM^1-105^-YFP marked as above. **(E-F)** Fluorescence intensity profiles across Pm for SP-YFP (E) and AvrM^1-105^-YFP (F). **(G)** Immunoblot analysis of flax leaf samples infected with transgenic *M. lini* strains as indicated. The arrows indicate bands at the expected molecular weight of AvrM^1-105^-YFP or SP-YFP. Protein loading is indicated by Ponceau staining (PS) of the Rubisco large subunit.

**Table 1.** Transformation gene constructs and parental and transformed *M. lini* strains. Construct Parental strains Transgenic strains Using the *AvrM* promoter and terminator.

| Construct | Parental strains | Transgenic strains |
| --- | --- | --- |
| Using the *AvrM* promoter and terminator |  |  |
| AvrM-YFP | CH5F2-96 | tAvrM-YFP-S1<br>tAvrM-YFP-S2 |
| SP-YFP | CH5F2-89 | tSP-YFP |
| AvrM<sup>1-105</sup>-YFP | CH5F2-89 | tAvrM<sup>1-105</sup>-YFP-S1<br>tAvrM<sup>1-105</sup>-YFP-S2 |
| AvrM<sup>1-156</sup>-YFP | CH5F2-89 | None recovered |
| CFP | CH5F2-89 | tCFP |
| Using the *AvrP123* promoter and terminator |  |  |
| AvrP123-YFP | CH5F2-74 | tAvrP123-YFP-S1<br>tAvrP123-YFP-S4 |

## Discussion

Molecular analysis of effector delivery from rust fungi has been hampered by the difficulties in genetic manipulation of these obligate biotrophs. In this study, we generated a set of transgenic *M. lini* strains expressing effectors as fusions with fluorescent proteins, allowing direct observation of effector localisation in live pathogen-infected host cells. *M. lini* transformation was achieved by *Agrobacterium*-mediated T-DNA delivery, using silencing of the *M. lini* effector gene *AvrL567* to select transformants by their virulence on flax lines containing the resistance gene *L6* (Lawrence *et al*., 2010). We were successful in transforming five of six constructs into *M. lini* using this approach. This enabled the first visualisation of the accumulation and localisation of effector-YFP fusions by fluorescence microscopy after delivery by *M. lini*.

The transgenic *M. lini* strains generated here express variants of the *AvrM* or *AvrP123* genes under the transcriptional control of their respective native promoters and terminators to maintain their normal expression patterns. Previous expression analysis indicated that *M. lini Avr* genes are expressed only during host infection and their transcripts are highly enriched in haustoria (Dodds *et al*., 2004; Catanzariti *et al*., 2006; Wu *et al*., 2019). In this study, all transgenically-expressed YFP fusion proteins (and CFP alone) driven by the native *AvrM* or *AvrP123* regulatory sequences accumulated in haustoria and to a lesser extent in haustorial mother cells, but were not found in infectious hyphae or substomatal vesicles or in the plant apoplast. This suggests that their expression is highly specific to haustoria and haustorial mother cells and does not occur in other cell types present during infection. Accumulation of YFP fluorescence was first detected four days after infection although transcripts could be detected as early as two dpi (Wu *et al*., 2019).

Both AvrM and AvrP123 effectors are delivered into the host cytsosol as evidenced by their recognition by host intracellular NLR proteins (Dodds *et al*., 2004; Catanzariti *et al*., 2006; Barrett *et al*., 2009; Anderson *et al*., 2016). Fluorescence and electron microscopy coupled with immunolocalisation previously detected the AvrM protein in both the EHMx and within infected host cells during infection (Rafiqi *et al*., 2010). Here, we found that intact AvrM-YFP and AvrP123-YFP fusion proteins both accumulated in the EHMx, consistent with their secretion from haustoria. AvrM-YFP was also observed within host cells, suggesting its subsequent transfer from the EHMx. AvrM-YFP was excluded from the host nucleus, also consistent with previous immunolocalisation data for infected tissue, as well as *in planta* transient expression data (Ve *et al*., 2013). The *M. lini* transgenic lines expressing AvrM-YFP were unable to infect flax lines containing the corresponding *M* resistance gene, confirming complementation of the avirulence phenotype by this fusion protein.

In contrast to AvrM, expression of AvrP123-YFP from transgenic *M. lini* resulted in the detection of fluorescence only in the EHMx but not within host cells. Furthermore, these transgenic lines were fully virulent on flax lines containing the corresponding resistance gene *P2*. We previously showed that AvrP123-YFP was recognised by P2 when expressed transiently *in planta* via *Agrobacterium*, suggesting that the YFP fusion does not disrupt the overall protein structure or its recognition (Zhang *et al*., 2018). However, fusion of YFP to this protein appears to prevent its delivery from the EHMx into host cells, although this was not the case for the AvrM-YFP fusion. It is possible that a signal region within the AvrP123 protein required for its delivery is obscured by the YFP protein. Using a longer peptide linker between the effector and YFP can be considered in the future to alleviate this problem. Another possibility is that the two effector proteins are delivered by different mechanisms, with only AvrP123 delivery incompatible with the YFP fusion. Notably, AvrM-YFP accumulated in bright structures within the haustorial body which may be nuclear associated. The nature and role of these fluorescence foci is not known.

We further dissected the AvrM sequence to identify regions involved in protein delivery into the host cell. The AvrM protein consist of an N-terminal 28 aa signal peptide (SP) and a C-terminal region (aa 106-343) that forms a stable dimer and interacts directly with the M protein, separated by an intervening region (aa 29-105) of unknown function (Catanzariti *et al*., 2010; Ve *et al*., 2013). We showed that the N-terminal 1-34 amino acids of AvrM containing the signal peptide are sufficient for secretion of YFP into the EHMx, but no fluorescence was detected in host cells. This suggests that delivery from the EHMx to the host cell requires a specific signal within the AvrM protein that is not present in YFP. In contrast, fusion of the first 105 amino acids of AvrM to YFP resulted in fluorescence in both the EHMx and within host cells, similar to full length AvrM-YFP. Unlike full-length AvrM-YFP, AvrM^1-105^-YFP accumulated in the host nucleus, probably due to its smaller size allowing diffusion through the nuclear pore. Thus, the region of AvrM between amino acids 34 and 105 appears to contain a separate signal directing delivery of the secreted protein into host cells. This further implies the existence of a delivery mechanism that recognises specific effector proteins for translocation into host cells, while others may be secreted and remain in the EHMx as observed for SP-YFP. These observations are consistent with a two-step process of AvrM delivery; first, secretion via a conventional SP-mediated pathway, followed by translocation into host cells from the EHMx as proposed for *U. maydis* (Ludwig *et al*., 2021). EHMx-resident proteins carrying a host-delivery signal may be substrates for a translocon present in the EHMx. Alternatively, it is also possible that recognition of the delivery signal occurs within the secretory pathway of the fungus, possibly co-incident with SP recognition and translocation, followed by diversion into a specialised delivery-specific pathway as suggested for oomycetes and *Magnaporthe* (Giraldo *et al*., 2013).

Previous analysis of AvrM proteins transiently expressed *in planta* found that these secreted proteins accumulated inside plant cells, despite the presence of a functional SP (Rafiqi *et al*., 2010). This intracellular accumulation was dependent on protein sequences in the region between amino acids 106 and 156 in AvrM and was not observed for the AvrM^1-105^ fragment, which accumulated in the plant apoplast. These observations had suggested the possibility of a pathogen-independent uptake mechanism into plant cells for rust effector proteins. Although similar observations were made for other fungal and oomycete effector proteins, these pathogen-free assay systems may produce artefactual results that are not reflective of effector delivery processes that occur during infection (Petre & Kamoun, 2014). Indeed, our results here indicate that the same AvrM^1-105^ fragment that was apoplastic when expressed *in planta* was correctly delivered into the host cytoplasm during fungal infection. These data suggest that the intracellular accumulation of *in planta* secreted AvrM does not fully reflect the rust effector delivery mechanism and highlight the importance of using a native rust pathogen-host system to study effector delivery. It will be important in future to establish whether the host CME pathway is involved in rust effector delivery during infection as has now been shown for *P. infestans* (Wang *et al*., 2023b) and *M. oryzae* (Oliveira-Garcia *et al*., 2023).

## Materials and Methods

### Plant and rust materials

The flax (*L. usitatissimum*) cultivars (*cv*.) Hoshangabad, Bison, Birio (containing the *R* gene *L6*), Dakota (*M*) and Abyssinian (*P2*), as well as near isogenic Hoshangabad or Bison backcross derivatives containing these resistance genes, have been described previously (Lawrence *et al*., 1981; Islam & Mayo, 1990). The flax *cv*. Hoshangabad does not contain any known *R* genes (Mayo & Shepherd, 1980) and was used for generation of *M. lini* transgenic lines. The *M. lini* rust strains CH5F2-89, CH5F2-96 and CH5F2-74 are homozygous for *AvrL567*, which confers avirulence on flax lines containing the corresponding *L5*, *L6* or *L7* resistance genes (Lawrence *et al*., 1981; Dodds *et al*., 2004). In addition, CH5F2-89 and CH5F2-96 lack *AvrM* and are therefore virulent on flax lines containing the *M* resistance gene, while CH5F2-74 lacks *AvrP123* and is virulent on flax lines containing *P2*.

### Vectors and gene constructs

The pTNsiAvrL567 vector encoding a hairpin construct that silences *AvrL567* has been described previously (Lawrence *et al*., 2010). A 1.1 kb fragment of the *AvrM* terminator region downstream of the stop codon was amplified from the *AvrM-A* genomic clone with XbaI-NcoI on the 5’ end and SbfI-SalI on the 3’ end and cloned into the XbaI and SalI sites of the pBluescript vector. The coding sequence of the yellow fluorescent protein (YFP) was inserted into the XbaI/NcoI sites and the coding region of *AvrM* including a 1.4 kb promoter region was amplified with SacI-SbfI and XbaI linkers and inserted into the SacI and XbaI sites of the vector. The AvrM and YFP coding sequences were fused via a gene linker encoding a GPGP (glycine-proline-glycine-proline) peptide. The full SbfI AvrM-YFP fragment including the *AvrM* promotor and terminator was then cut from pBluescript and cloned into the PstI sites of the pTNsiAvrL567 vector, resulting in replacement of the spectinomycin resistance gene cassette downstream of the AvrL567 silencing construct (Fig S2). The AvrM N-terminal constructs including SP-YFP were generated based on pBluescript or pTopo AvrM-YFP using site-directed mutagenesis with corresponding primer sets and cloned into pTNsiAvrL567 as described above (Table 1). A construct containing the cerulean variant of cyan fluorescent protein (CFP) was made without any signal peptide and inserted between the AvrM promotor and terminator (Table 1).

The AvrP123 promoter region consisting of 1.7 kb upstream of the start codon (Barrett *et al*., 2009; Nemri *et al*., 2014) was amplified from rust genomic DNA with a KpnI-SbfI linker at the 5’ end and a blunt 3’ end, and cloned into the KpnI and EcoRV sites of pBluescript, generating pBScAvrP123-pro. The gene encoding AvrP123-YFP was amplified from a pNos-AvrP123-YFP vector (Zhang *et al*., 2018) as a blunt (5’) and XmaI (3’) fragment and cloned into the EcoRV and XmaI sites of pBScAvrP123-pro vector, generating pBScpro-AvrP123-YFP. The AvrP123 and YFP coding sequences were linked via a glycine-proline-glycine-proline (GPGP) peptide. An 800 bp AvrP123 terminator region was amplified from the *M. lini* genome and cloned into the SmaI site of pBScpro-AvrP123-YFP as a blunt fragment, resulting in a SbfI site at the 3’ end. The full SbfI AvrP123-YFP was cloned into the PstI sites of the pTNsiAvrL567 vector.

The mCherry variant of the red fluorescent protein (RFP) used for transient expression *in planta* was assembled using the Plant Molecular Golden Gate Cloning kit (Engler *et al*., 2014). The RFP coding gene was inserted in between the CaMV 35S promoter and terminator and cloned into the pICH47732 binary vector for *Agrobacterium* transformation.

### *M. lini* transformation and inoculation

*M. lini* strains CH5F2-96, CH5F2-89 or CH5F2-74 were transformed using the *Agrobacterium*-mediated method described by Lawrence *et al*. (Lawrence *et al*., 2010). In brief, the parental rust strains were inoculated onto flax *cv*. Hoshangabad and *Agrobacterium* strains carrying each transformation construct were introduced into the stems of the flax plants 5 days post inoculation (dpi). The plants were kept in a glasshouse and urediospores were collected after 9 days and at subsequent intervals of 4 or 5 days. The spores were then inoculated onto flax lines carrying *L6* to select transformants. Single pustules of transgenic rust strains were collected and increased on *cv*. Hoshangabad flax plants.

For phenotyping and microscopic studies, *M. lini* urediospore stocks were resuspended in Novec 7100 (3M) in a 1/10 ratio spore/fluid (v/v) and applied to the leaves of 4-week old flax plants using a paintbrush. The inoculated plants were sprayed with a fine mist of water and kept in high humidity at 20°C overnight in a plastic bin. The plants were then moved to a growth cabinet maintained at 24°C with a 12/12 h light-dark cycle.

Flax plants *cv*. Hoshangabad were inoculated with transgenic *M. lini* strains expressing various AvrM-YFP fusion constructs prior to labelling the cytosol of flax cells by transient expression of RFP. An Agrobacterium strain containing the 35S::RFP construct was cultured in LB media at 28°C overnight and harvested by centrifugation. The cells were resuspended to an OD_600_ of 1 in an infiltration buffer containing 10 mM MES pH 5.6, 10 mM MgCl_2_ and 200 µM acetosyringone. The Agrobacterium resuspensions were introduced to infected flax leaves using a syringe at 3 dpi. Leaf samples were collected and imaged at 5 dpi.

### Quantification of fungal biomass by chitin measurement

Fungal biomass was quantified using the wheat germ agglutinin chitin assay (WAC) as described by Ayliffe *et al* (Ayliffe *et al*., 2013). In brief, flax leaf tissue with or without *M. lini* strains was harvested 10-11 dpi and snap-frozen in liquid nitrogen. Each sample contains two 2.5-cm flax leaves. The tissues were autoclaved in 1 M KOH containing 0.1% (vol/vol) Silwet L-77 and neutralized in 50 mM Tris, pH 7.0. The tissues were then homogenised using a tissuelyser (Qiagen). A 100-µl sample of each tissue suspension was added to 2 µl of a 1-mg/ml solution of the lectin wheat germ agglutinin (WGA) conjugated to fluorescein isothiocyanate (FITC) (Sigma Aldrich) and incubated for 30 min at room temperature. The samples were washed three times, resuspend in 150 µl 50 mM Tris (pH 7.0) and 150 µl of each sample was transferred to a black 96-well plate. Fluorometric measurements were made using a plate reader (Tecan) with 485-nm excitation wavelength and 535-nm emission wavelength.

### Protein extraction and immunoblotting

Total proteins were extracted from *M. lini*-infected flax leaves collected 7 dpi by grinding in Laemmli buffer (Laemmli, 1970). Proteins were separated by SDS-PAGE and transferred to nitrocellulose membranes (Bio-Rad). Membranes were blocked using 5% (w/v) skim milk in TBST buffer [50 mM Tris-HCl Ph 7.5, 150 mM NaCl and 0.1% (v/v) Tween 20] and probed with anti-GFP mouse monoclonal antibodies (Roche) followed by goat anti-mouse horseradish peroxidase (HRP) conjugate (BioRad). Immunolabelling of the proteins was detected with the SuperSignal West Femto chemiluminescence kit (Thermo Scientific). Membranes were stained with Ponceau S to visualize protein loading.

### Confocal microscopy

Freshly cut flax leaf samples were mounted in water between glass slides and viewed on a Zeiss LSM 780 confocal microscope (Carl Zeiss Microscopy) with an LD C-Apochromat 40x/1.1 W Korr M27 water immersion objective. Images were collected using an excitation wavelength of 514 nm and a collection window of 516–596 nm for YFP and 650–720 nm for chlorophyll autofluorescence. The RFP fluorescence was imaged with an excitation wavelength of 561 nm and a collection window of 579-650 nm. To separate YFP from multiple fluorophores with a high degree of overlap in their emission spectra, spectral data were acquired using the 18-channel lambda-mode in the ZEN software (Carl Zeiss Microscopy), with 514 nm excitation and a collection window of 517-650 nm. The spectral data were processed using the linear unmixing function in ZEN, and the resulting YFP profiles were compared with the manufacturer’s data to confirm their identity. The fluorescence intensity values across a linear region of interest (ROI) were measured using the Fiji package (Schindelin *et al*., 2012). The results were plotted and graphed in Microsoft Excel.

## Acknowledgments

We thank Daryl Webb and Leila Blackman for advice and technical assistance on confocal microscopy, and the Centre for Advanced Microscopy and Microscopy Australia for providing microscopic facilities.

## Author Contributions

JPR, DAJ, and PND conceptualized the project, acquired funding and supervised the work. XZ, AC, GJL and PHPG acquired experimental data and conducted data analysis. XZ drafted the manuscript and all authors contributed to writing, reviewing and editing.

## Data Availability

The data that support the findings of this study are available from the corresponding authors upon reasonable request.

## Supplementary figure legends

**Fig S1. Fluorescence emission spectra showing four fluorophores (ACE1-ACE4) identified by linear unmixing.**

The YFP reference spectra are added and shown as yellow dashed lines. The confocal microscope images were taken at 3 dpi **(A)** and 4 dpi **(B)**. YFP fluorescence is shown as a yellow line. Chloroplast autofluorescence is shown in red; undefined fluorescence emissions are shown in blue and cyan.

**Fig S2. Constructs for *M. lini* transformation.**

Schematic representation (not to scale) of the T-DNA constructs used to transform *M. lini*. The T-DNA contains the *AvrL567* silencing construct (siAvrL567) between the left (LB) and right (RB) borders. The siAvrL567 construct expresses an inverted repeat *AvrL567* transcript driven by the *AvrL567* promoter (prom). Genes encoding *AvrM* or *AvrP123* (Avr)-YFP fluorescent proteins were inserted into the *Pst*I restriction site between siAvrL567 and the RB. The *Avr-YFP* genes are under the control of the native *AvrM* or *AvrP123* promoter (Avr prom) and terminator (Avr term) sequences. The internal *Pst*I restriction sites are indicated above the cartoon.

