## Supplementary figures and images for "Translocation of effector proteins into plant cells by the flax rust pathogen *Melampsora lini*"

### Fig S1

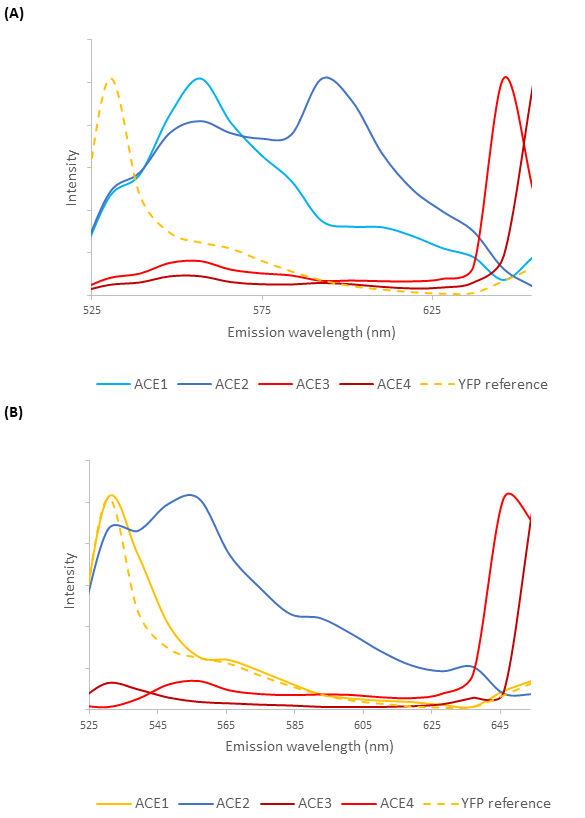

### Fig S2

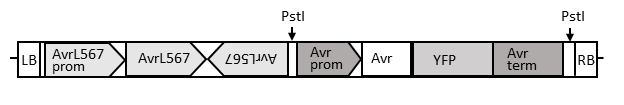
